# Architecture of the spinach plastid-encoded RNA polymerase

**DOI:** 10.1101/2024.02.16.580782

**Authors:** Tongtong Wang, Guang-Lei Wang, Ying Fang, Yi Zhang, Wenxin Peng, Yue Zhou, Aihong Zhang, Long-Jiang Yu, Congming Lu

## Abstract

The plastid-encoded RNA polymerase (PEP) serves as the principal transcription machinery within chloroplasts and is responsible for transcribing over 80% of primary plastid transcripts. Plant PEP is composed of a prokaryotic-like multisubunit core enzyme known as the PEP core, supplemented by newly evolved PEP-associated proteins (PAPs). In this study, we present the cryo-electron microscopy (cryo-EM) structure of a PEP complex derived from spinach (*Spinacia oleracea*). Our structural analysis unveils the presence of 14 PAPs closely associated with the PEP core complex consisting of 5 subunits, forming an irregular-shaped supercomplex. We provide a detailed depiction of the PEP core subunits and the 14 PAPs, elucidating their precise spatial arrangement and interactions. This research offers a crucial structural basis for future investigations into the functions and regulatory mechanisms governing plastid transcription.

---

Chloroplasts are the photosynthetic organelles in eukaryotic organisms, and play a crucial role in capturing sunlight and converting it into energy. These organelles possess their own genome, which is a relic of an endosymbiotic event. The chloroplast genome is relatively small, encoding only 75–80 proteins out of the approximately 3500–4000 proteins present in chloroplasts^1,2^. However, the proper expression of these genes is vital for chloroplast biogenesis, as well as the growth and development of plants^3,4^.

Transcription of chloroplast genes is carried out by two distinct types of RNA polymerases: the nuclear-encoded polymerase (NEP), resembling the T3-T7 phage-type RNA polymerase, and the plastid-encoded polymerase (PEP), which is a bacterial-type multisubunit polymerase^5,6^. PEP represents the primary transcription machinery in chloroplasts and predominantly transcribes genes related to photosynthesis, generating the majority of mRNA, tRNA, and rRNA molecules^7,8^.

In land plants, PEP exhibits a bacterial-like RNA polymerase with a catalytic core inherited from its cyanobacterial ancestor. This core consists of α, β, β’, and β’’ subunits encoded by the *rpoA, rpoB, rpoC1*, and *rpoC2* genes, respectively, located in the plastid genome^4,9^. Knockout mutants in tobacco plants deficient in any of these *rpo* genes exhibit albino or yellowish phenotypes and seedling lethality due to impaired chloroplast development, underscoring the indispensable role of Rpo subunits in chloroplast biogenesis. Additionally, the PEP core relies on nuclear-encoded σ factors for promoter recognition and transcription initiation^10^.

Despite genetic evidence pointing toward a prokaryotic structure of the PEP core^9,11^, biochemical purification of PEP from plant chloroplasts has revealed a much higher number of subunits^12^. In *Arabidopsis*, at least 12 distinct PEP-associated proteins (PAP1-PAP12) have been identified as tightly associated with the PEP core^4,9^. Genetic studies have also demonstrated that the loss of these PAP proteins leads to severely impaired PEP-mediated transcription and albino or yellowish phenotypes, highlighting the essential nature of these PAPs for PEP activity and revealing the intricate organization of PEP^13-20^. However, the specific roles of individual PAPs remain poorly understood. Notably, PAP2/pTAC2, PAP3/pTAC10, PAP5/pTAC12, and PAP11/MurE are crucial for the assembly of a functional PEP complex, suggesting that PAPs are essential for establishing the structural foundation of PEP transcriptional activity^21,22^.

Despite these significant insights, the exact structural basis of the molecular functions of the PEP transcription machinery has remained elusive. In this study, we present the cryo-EM structure of the PEP complex isolated from spinach, unveiling intricate structural features that shed light on the mechanism of PEP-mediated transcription in higher plants.

## Overall structure

The PEP complex was isolated and purified from spinach (Extended Data Fig. 1), and its structure was determined by cryo-EM at an overall resolution of 3.16 Å (Fig.1, Extended Data Fig. 2 and Extended Data Table 1), the obtained maps were of sufficient quality to allow building and refinement of an almost complete model of the PEP complex with the assistance of AlphaFold2^23,24^, revealing the features of the PEP complex at molecular detail. The whole PEP complex displayed an obviously asymmetric structural conformation with a total molecular weight of approximately 960 kDa (Extended Data Fig. 1). We identified totally19 subunits in the whole PEP complex (Fig.1, Extended Data Fig. 3), among which 14 subunits were reported previously^4,12^, including α, β, β’, β’’, PAP1 and PAP3–PAP12. Additional two subunits, FLN2 and pTAC18 were identified based on the electron density map. Among which, FLN2 contains a pfkB-type carbohydrate kinase domain and shares structural similarity with PAP6/FLN1 (Extended Data Fig. 4a), while pTAC18 lacks clear structural domains (Extended Data Fig. 3). We designated FLN2 and pTAC18 as PAP13 and PAP14, respectively.

**Figure 1.**
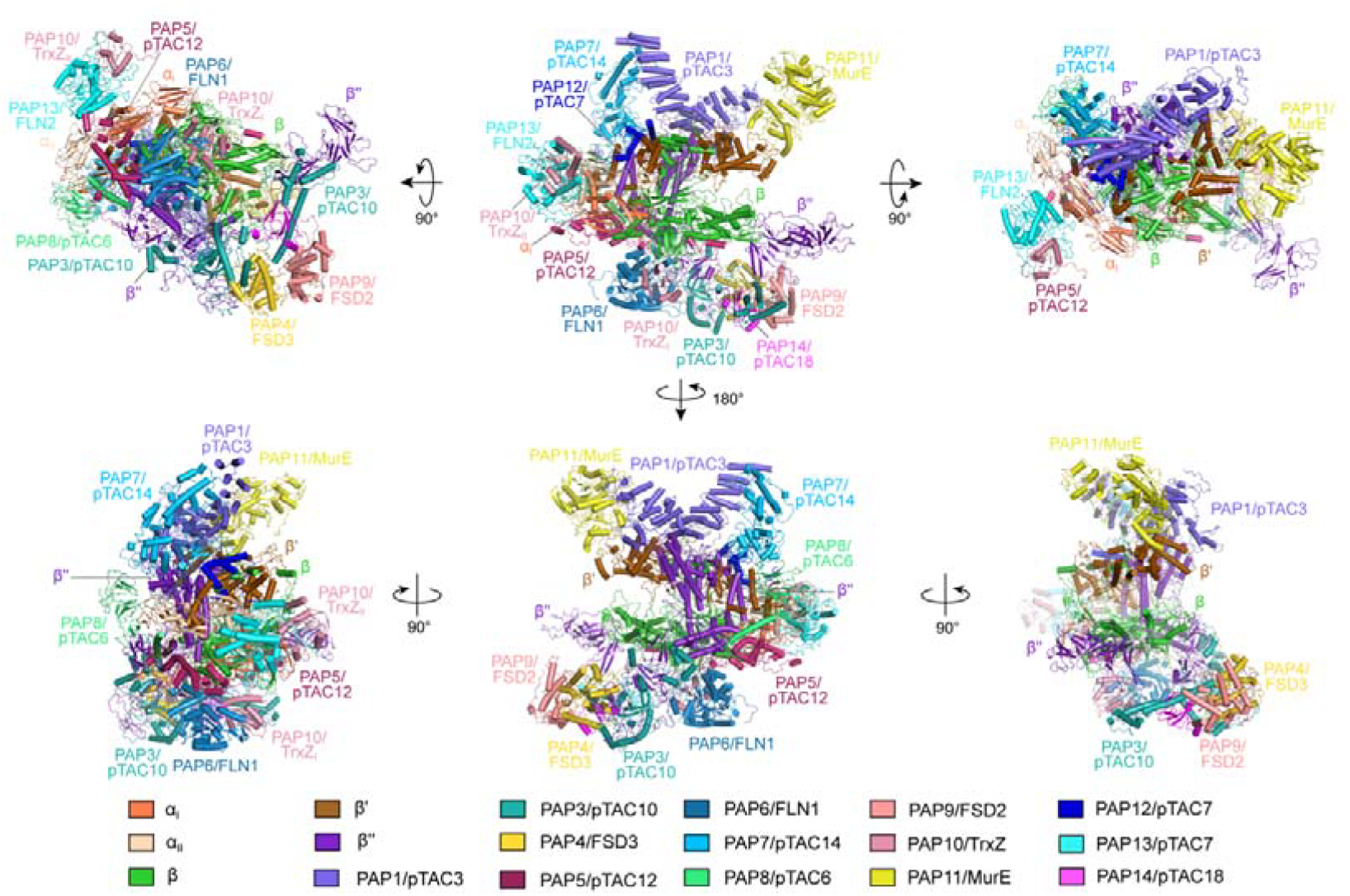
Overall architecture of the spinach PEP complex. Different views show the overall arrangement of the PEP subunits. Protein subunits are shown as cartoons and colored individually as indicated.

Another previously reported PAP2 was detected weakly in the purified PEP complex by SDS-PAGE electrophoresis (Extended Data Fig. 1d), however, it was not identified in the present structure, suggesting that PAP2 associates loosely in the PEP complex and may be lost for the structure determination. Additionally, some fragmented electron densities were not continuous in the cavity formed by α, β, PAP5 and PAP6 subunits (Extended Data Fig. 5a) and considered to be the component of β’’, since it has a long loop region based on the prediction of AlphaFold2 (Extended Data Fig. 5b) and these regions could not be traced well in this study.

Within the PEP complex, there is an essential catalytic core that is composed of five subunits (α_2_ββ’β’’), and it is encircled by 14 PAP protein subunits, forming a structure resembling a “crab claw” with two arms, featuring an internal groove or channel along its length. From the front view (Fig.1 upper middle), we could observe the classic crab claw structure, including two arms: the upper arm and the lower arm. PAP1, PAP7, and PAP11 locate above the upper arm. PAP6 and PAP10 are positioned just below the lower arm, while PAP3, PAP4, PAP9, and PAP14 locate at the back of the lower arm. PAP5, PAP8, PAP13, and PAP10 are located at the intersection of the two arms. Interestingly, two copies of PAP10 interacted with PAP6 and PAP13 to form a heterodimer, respectively, were identified and located at different positions in the structure.

## Structure of the PEP core complex

Chloroplasts originated approximately 1.5 billion years ago from ancient cyanobacteria that were engulfed by a eukaryotic host cell. The RNA polymerase (RNAP) of cyanobacteria consists of 6 subunits (α_2_ββ’β’’ω), forming a structure resembling a “crab claw” with two arms^25,26^. The PEP core also shows a “crab claw” with two arms (the lower arm and the upper arm) (Fig. 2a), indicating that the overall structure of catalytic core from land plants owns a highly structural similarity with RNAP of cyanobacteria, despite relatively low sequence identity (Extended Fig. 6).

**Figure 2.**
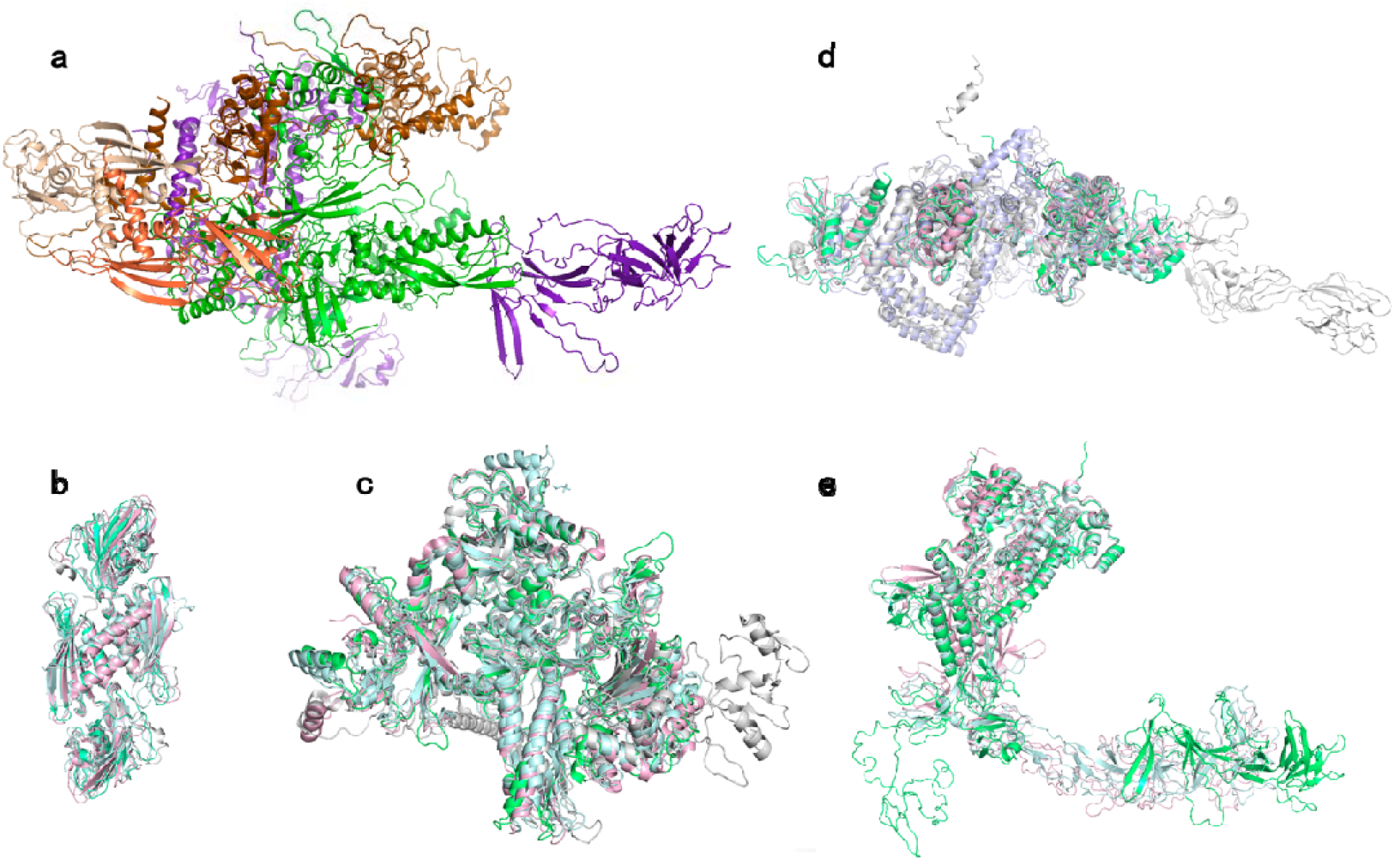
Structure of the spinach PEP core complex. Comparisons of the spinach PEP core complex with RNA polymerases from *Synechocystis sp*. PCC 6803 (PDB code: 8GZG; pale cyan), *Synechococcus elongatus* (PDB code: 8URW; light pink), *E. coli* (PDB code: 6GH5; light blue), *Thermus termophilus* (PDB code: 3DXJ; light gray) were made to show the major difference among these species. **a**, The spinach PEP core complex is composed of five subunits, including α_2_, β, β’, and β’’. All subunits are colored individually as indicated in Figure 1. **b, c, d and e** show the comparison of α_2_, β, β’, and β’’ subunits, respectively, In the comparison, the spinach PEP core complex is colored as limegreen.

The structural comparisons of the PEP core with RNAP from cyanobacteria and *E. coli* revealed that the structures of α dimer and β subunit are highly conserved in *E. coli*, cyanobacteria, and land plants (Fig. 2b, c). However, significant differences are observed in the structures of β’ and β’’ subunits between the PEP complex and bacterial RNAP (Fig. 2d, e). Unlike the PEP complex, the bacterial RNAP does not have β’’ subunit. The comparison of the structures of the β’ subunit between bacterial RNAP and the PEP complex shows that the whole structure of the β’ subunit in the PEP complex is well aligned with the part structures of the β’ subunit in bacterial RNAP (Extended Fig. 7a). We further made the structural comparison between the β’ subunit in bacterial RNAP and the β’’ subunit in the PEP complex (Extended Fig. 7b). We observed that the part structures of the β’ subunit in bacterial RNAP are highly superposed with the part structures of the β’’ subunit in the PEP complex (Extended Fig. 7b).

We also made structural comparisons of β’ and β’’ between the PEP complex and cyanobacterial RNAP. The structures of β’ subunit are highly conserved in cyanobacteria and plants (Extended Fig. 7c, d). The β’’ subunit of RNAP in cyanobacteria contains a Sequence Insertion 3 domain (SI3) that is located between the two helices of the Trigger Loop, a key mobile element in the catalytic center of RNAP. Previous structure shows that the SI3 domain in cyanobacteria is highly flexible and it regulates multiple stages of transcription by influencing the Trigger Loop folding^27^. The β’’ subunit of the PEP complex was also considered to contain a SI3 domain that is similar to that of cyanobacteria (Extended Fig.5b). However, due to the flexibility of β’’ subunit as predicted and the poor electron density around this region, it is impossible to generate a complete structural model of the SI3 domain (Extended Fig.5b) and the structure and function of the SI3 domain of PEP remain to be further studied.

During plant evolution, the gene encoding the ω subunit was considered to be lost from the plastid genome^9^. Intriguingly, the location and structure of PAP12 are similar to those of the ω subunit in cyanobacterial RNAP (Extended Fig. 8). Thus, we define PAP12 as ω subunit of the PEP complex and it play the same role with that of cyanobacterial RNAP in the transcription.

## Interactions between PAPs and the core complex

The PEP complex is a hybrid system comprised of a prokaryotic core and peripheral PAPs of eukaryotic origin^4^. Our structure reveals the composition and arrangements of these PAPs for the first time. In the overall structure, the locations of 14 PAP subunits can be divided into three regions, with two regions surrounding the two arms of the PEP core complex, forming a larger clamp-like structure.

Located on the upper arm are PAP1, PAP7, PAP11, and PAP12. PAP1 interacts with the β’ protrusion structure in close proximity to the PEP complex, forming interactions with the β, β’, and β’’ core subunits through a helical region (Extended Fig. 9a). The N-terminal structural domain of PAP1 contains seven helix-loop-helix motifs, forming a superhelical ribbon-like sheet, likely possessing nucleic acid binding activity (Extended Fig. 9a), which is consistent with the results that PAP1 binds to the promoter region of PEP-dependent genes^20^. In cyanobacterial RNAP, the β’ subunit associates with the σ factor responsible for bacterial RNAP and promoter recognition^27^. These findings suggest that PAP1 may be involved in the transcription initiation process of PEP. Although PAP2 was not identified in our structure, cross-linking mass spectrometry detected interactions between PAP1 and PAP2 in the PEP complex purified from *Arabidopsis*, implying that PAP1 and PAP2 may function together^28^. PAP11 specifically positioned at the forefront of the upper arm adjacent to the β’ subunit. Genetic and biochemical studies demonstrate that disruption of PAP11 leads to no accumulation of fully assembled PEP complex^22^. This suggests that PAP11 is essential for maintaining the structure of the PEP complex though it is positioned at the forefront of the upper arm. PAP7’s C-terminus interacts with the PAP1 protein, while its N-terminus interacts with PAP8.

Six PAPs (PAP3, PAP4, PAP6, PAP9, PAP10, PAP14) are situated in the lower arm (Fig. 3b). PAP6 and PAP10 locate closely adjacent to β subunit and they interact with each other through their antiparallel β-folds (Extended Fig. 9c), which is consistent with previous biochemical study showing that PAP6 interacts with PAP10^13^. PAP6 and PAP10 also interact with the β’’ and β’ subunit through their respective long loops. Our structure shows that PAP4 and PAP9 form a heterodimer as reported previously^17^, and it is adjacent to the β’’ subunit (Extended Fig. 9d). Both PAP4 and PAP9 interact with PAP3 through their flexible loops (Extended Fig. 9d).

**Figure 3.**
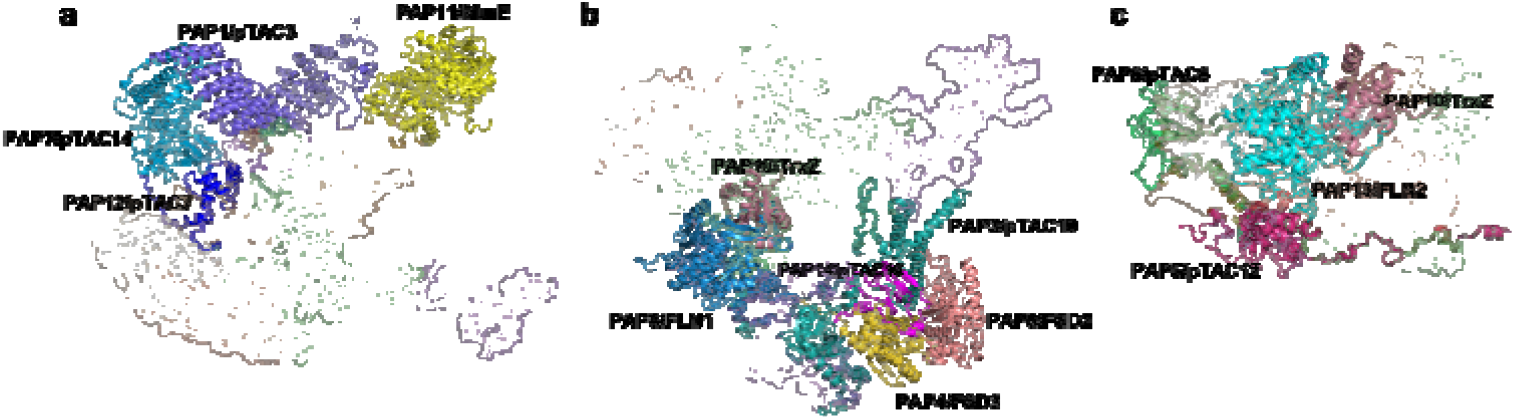
Arrangements of the 14 PAP subunits around the spinach PEP core. **a, b**, and **c** show the PAP subunits located on the upper arm, the lower arm, and the intersection of the two arms are shown, respectively. The structures of these PAPs are presented as cartoon, while the core complex as surface mode. All subunits are colored individually as indicated in Figure 1.

The PAP3 protein shows a long span with a length of about 131 Å. The C-terminal helix of PAP3 connects the PAP4 and PAP9 heterodimer and the β’’ subunit and interacts with PAP14 and the remaining part interacts with the irregular loops of the β’’ subunit (Extended Fig. 9e). The broad interactions of PAP3 with other PAPs and the β’’ subunit suggest that PAP3 is important for stabilizing the PEP complex, which is in line with the previous results showing that the *pap*3 mutant shows no accumulation of the fully assembled PEP complex and *OsPAP3* overexpression increases the expression of the PEP-dependent genes^14,22,29^.

PAP5, PAP8, PAP10, and PAP13 are positioned at the intersection of the two arms, near the α dimer (Fig. 3c). Biochemical and genetic studies have shown that disruption of PAP5 results in no accumulation of the fully assembled PEP complex and impaired transcription of the PEP-dependent photosynthesis genes^14,22^. Our structure shows that PAP5 protein, like a long thread with a lot of random loops, connects all subunits on the surface of the PEP core (Extended Fig. 9f). It is thus suggested that PAP5 is also a key component for stabilizing the PEP core complex. The C-terminus of PAP8 forms two short sheets that create an antiparallel β-fold within the β’ subunit, with helices extending into the region enclosed by an antiparallel β-fold and loop of the β’ subunit, forming an interacting area. PAP8 also interacts with β’ and β’’ through its long loop stretching into the PEP core and connects the upper arm through PAP7 (Extended Fig. 9b). PAP10 and PAP13 are adjacent to the α dimer.

## Structural insights into the molecular mechanism of PEP complex

Through a comparative analysis of the structures of the spinach PEP complex and the cyanobacterial RNAP, particularly focusing on the cyanobacterial RNAP in the initiation stage, these results provide a profound comprehension of the transcriptional machinery unique to chloroplasts^27^. Firstly, the PEP core bears a striking resemblance to that of cyanobacterial RNAP. Specifically, PEP exhibits a σ factor and DNA binding site analogous to those in cyanobacterial RNAP (Fig. 4a). Upon superimposition of these two structures, it becomes evident that none of the surrounding subunits clash with this binding site (Fig. 4b), indicating that PEP possesses a σ factor and DNA binding site akin to those in cyanobacterial RNAP. Secondly, within the cyanobacterial σ factor, there exists a loop region of approximately ∼50 amino acids in close proximity to the PAP11 subunit. In our structure, the electron density of PAP11 suggests a high degree of flexibility, indirectly implying a role for PAP11 in σ factor recognition or stabilization. Furthermore, in the presence of a σ factor and DNA binding, the SI3 head group of cyanobacterial β’’ subunit displays a stable conformation^27^, contrasting the highly disordered structure observed in our model. This strongly suggests that SI3 plays a pivotal role during the transcription process and undergoes significant conformational changes from open stage to the transcription initiation stage, shifting from a distal state to a closer site around the σ factor, forming an arch-shaped interaction (Fig. 4d). Lastly, at the DNA entrance site in spinach PEP complex, a small mass of electron density was observed, effectively obstructing the entrance. This position corresponds to the vicinity of PAP1, and the electron density is considered to represent a substantial missing loop region (∼200 amino acids) within PAP1. This observation implies an active role for PAP1 in processes associated with transcription, highlighting its potential involvement in the coordination and regulation of transcriptional events in chloroplasts.

**Figure 4.**
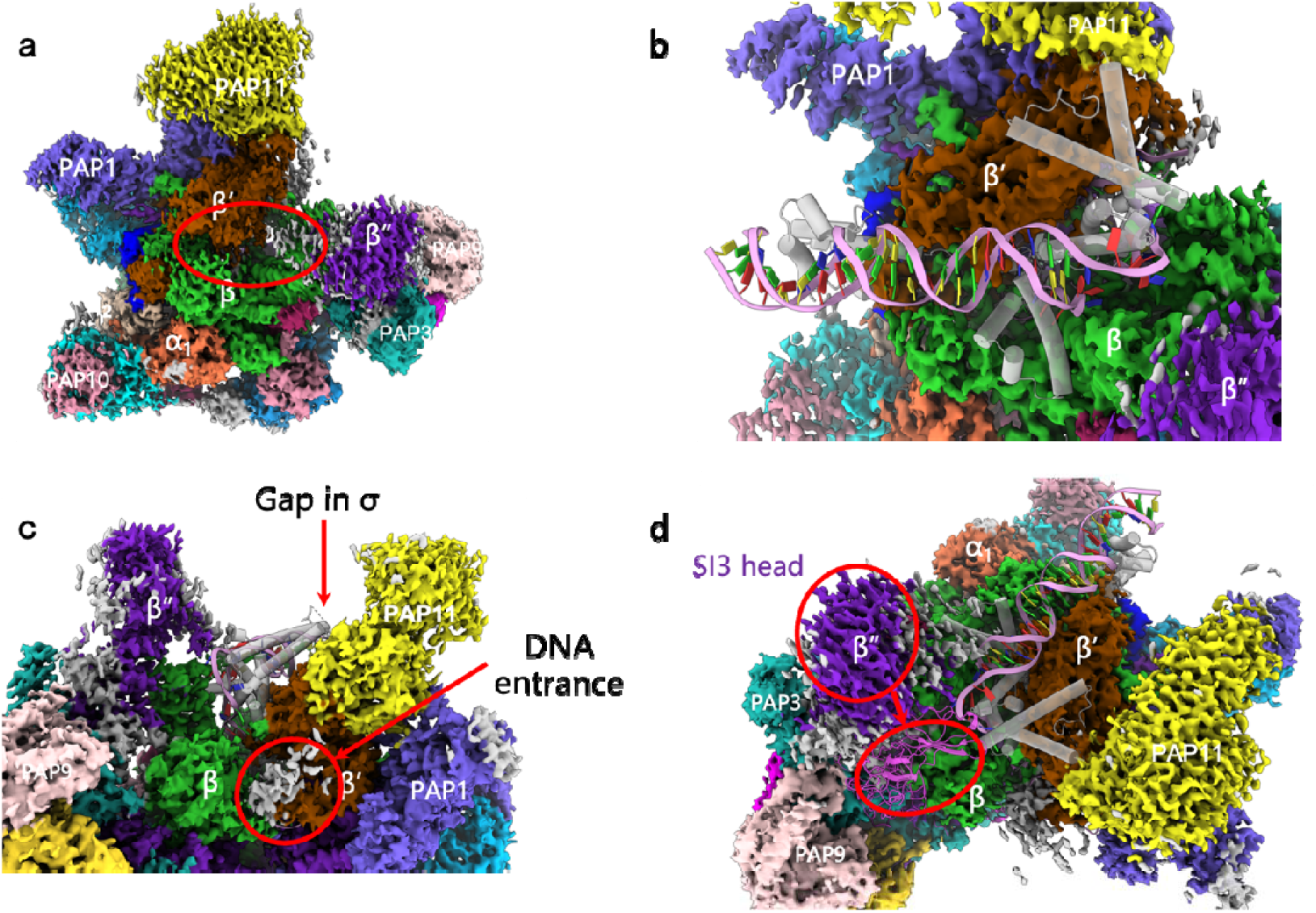
Proposed structure of the transcription initiation state of PEP complex. **a**, Density map of the PEP complex. The possible binding sites of DNA and σ cofactor are highlighted in red circle. **b**, Proposed structure of PEP-σ-DNA complex. The model is constructed based on the comparison of PEP complex and RNAP from cyanobacteria. DNA is shown in a pink ribbon, and the σ cofactor is shown as a gray cartoon model. **c**, Lateral view of the σ binding site. **d**, Potential conformational changes of the SI3 domain between “open” state (shown as density map) and σ-binding state (shown as purple cartoon). The models of σ and DNA are sourced from structure PDB: 8GZH.

In summary, we present here the structure of spinach PEP complex at “open” state which means there are no DNA and σ factor existed in the complex. The structure reveals that PAPs play a crucial role in sustaining the activity of the PEP complex by facilitating the assembly of a fully functional PEP complex as demonstrated previously^22^ and it can be inferred that PAPs are indispensable for establishing the structural integrity of the PEP complex to fulfill the function. The structural comparisons between PEP and cyanobacterial RNAP, particularly in the context of cyanobacterial RNAP’s initiation stage, provide valuable insights into the molecular mechanisms of PEP and highlight the potential roles of various subunits in transcription processes. It is plausible to suggest that PAPs might have been acquired during the course of plant evolution, possibly serving to uphold and potentially enhance the transcription of genes related to photosynthesis in higher plants.

## Limitation of the present study and some perspectives

In our present structure, a large flexible loop region of β’’ subunit (residue 562-862) predicated by AlphaFold2^23,24^ was located in the outer space of the PEP complex with a different conformation when compared with all known structures of RNAP (Fig. 2, Extended Fig. 5b), although these residues could not be traced ambiguously. This loop region was proposed to act as a regulatory valve in the different transcription stage. In addition, one of the most important PAPs, pTAC2 was also not identified in the structure, suggesting a potential loose association with core complex or peripheral subunits. The inability to resolve these detailed structures is an obvious indicator of their function, as dynamic conformational changes would be needed for their role in DNA recognition and RNA synthesis. Future structural and functional studies are required to further elucidate the mechanisms for chloroplast RNA polymerase recognition and specificity, and its discrimination from related factors. Additionally, our structure did not include its corresponding substrate DNA and σ factor, which is believed to stabilize the PEP complex at different stages of transcription, such as initiation, pausing, elongation and termination. The nature and mode of action of the PEP complex still requires further investigations.

## Methods

### Data reporting

No statistical method was used to predetermine the sample size. The experiments were not randomized, and the investigators were not blinded to allocation during experiments and outcome assessment.

### Purification of the spinach PEP complex

About 5 kg of mature spinach leaves were cut into small pieces. Leave pieces were gently homogenized by a blender in 15 L of pre-cooled Chloroplast Isolation Buffer (CIB, 20 mM HEPES, pH 8.0, 0.33 M sorbitol, 5 mM MgCl_2_, 5 mM EGTA, 5 mM EDTA, 10 mM NaHCO_3_, 60 mM sodium ascorbate). The resulting homogenate was filtered through three layers of Miracloth (Millipore). The crude chloroplasts were collected by centrifugation at 3,000 g for 5 min at 4ºC and further lysed with 2 L of Chloroplast Breaking Buffer (50 mM HEPES, pH 7.6, 4 mM EDTA, 20% (vol/vol) glycerol, 20 mM β-mercaptoethanol, 50 μg ml^−1^ phenylmethylsulfonyl fluorid (PMSF), 0.1 M (NH_4_)_2_SO_4_). The insolubilized materials were removed by centrifugation at 40,000 g for 30 min at 4ºC. The supernatant was applied to a Heparin Sepharose CL-6B column (1.6 cm diameter, 20 cm length, column volume 31 ml). The column was washed with 100 ml buffer containing 50 mM HEPES, pH 7.6, 4 mM EDTA, 10% (vol/vol) glycerol, 20 mM β-mercaptoethanol, 50 μg ml^−1^ PMSF. Bound proteins in the column were eluted with 60 ml of 0.28 M (NH4)_2_SO_4_, and further separated by a 0.44–0.88 M continuous sucrose density gradient in a buffer (50 mM HEPES, pH 7.6, 0.1 mM EDTA, 1 mM DTT, 0.5 × proteinase inhibitor cocktail, and 5% (vol/vol) glycerol) using a Beckman Coulter Optima XPN-100 centrifuge with SW40Ti rotor at 16,0000 g at 4ºC for 23 h. After centrifugation, the 6.5 to 9 ml sample fractions from the top of sucrose density gradient were collected and concentrated to a concentration of 5 mg ml^−1^ using a membrane concentrator with a 100-kDa cut-off. Then, the sample were subjected to gel filtration chromatography (Cytiva, Superose 6 Increase 10/300GL) in buffer containing 50 mM HEPES, pH 7.6, 2 mM EDTA, 2% (vol/vol) glycerol, 10 mM β-mercaptoethanol, and 50 μg ml^−1^ PMSF. The peak fraction from plastid-encoded RNA polymerase complex was collected for cryo-EM analysis.

### Characterization of the spinach PEP complex

The protein compositions of the purified PEP complex were analyzed by polyacrylamide gel electrophoresis (SDS–PAGE) using a gel containing 15% (wt/vol) polyacrylamide. The gels were stained with Coomassie brilliant blue (CBB) R-250. For mass spectrometry (MS) analysis, CBB-stained bands were cut out from the gel, destained in 25 mM NH_4_HCO_3_/50% (vol/vol) acetonitrile, alkylated with 25 mM iodoacetamide at 25°C in the dark, and digested with trypsin 37°C. The resulting peptides by sonication in buffer containing 5% (vol/vol) trifluoroacetic acid and 50% (vol/vol) acetonitrile, separated by reverse-phase liquid chromatography with a 150 μm × 250 mm analytical column packed with C18 particles (1.9 µm diameter), and measured in an Orbitrap analyzer at 240,000 resolution (at 400 m/z) with a target value of 106 ions. The tandem mass spectrometry (MS/MS) spectra were searched against *Spinacia oleracea* protein sequences downloaded from UniProt using MaxQuant (version 1.6).

### Cryo-EM data collection

The concentration of the purified PEP complex was adjusted to about 2 mg mL^-1^. Three microliters of protein solution were applied to a glow-discharged (Solarus plasma cleaner, Gatan) holey carbon grid (Quantifoil grid R2/1, 200 mesh, Cu) that had been treated with H_2_ and O_2_ mixtures for 30 s. The grid was plunged into liquid ethane cooled by liquid nitrogen using Vitrobot Mark IV (Thermo Fisher Scientific). The parameters were set as follows: blot force 0, blotting time 3 s, humidity 100%, sample chamber temperature 8°C. Data were collected on a Titan Krios (Thermo Fisher Scientific, Hillsboro, USA) electron microscope at 300 kV equipped with a K3 camera (Gatan) and at nominal magnification of 62 k in super-resolution mode. Movies were recorded in super-resolution mode and Fourier-cropped to give a resulting calibrated pixel size of 1.10 Å at the specimen level. An exposure rate of 22.5 e^-^ per pixel per s was set and a fresh super-resolution gain reference was performed at this dose rate before data acquisition. A total dose of 50 e^-^ per Å^2^ was used for movies of 40 frames. In total, 7,175 movies for the PEP complex were collected with defocus values from –0.8 to –2.0 μm.

### Image processing of the spinach PEP complex

All frames Motion Correction and CTF Estimation were performed with CryoSPARC ^30^. F-crop factor was set to 1/2 to generate aligned micrographs in Motion Correction. Particles were picked by crYOLO^31^ using a pretrained model. A total of 2,338,580 particles were picked from 7,175 micrographs at 0.01 threshold and 200 box size. These particles were then imported to CryoSPARC at 512-pixel box size, and downsampled to 128-pixel boxsize for 2D classification with 100 classes. 1,266,268 particles were selected for a secondary 2D classification with 50 classes. 384,187 particles from 21 good classes were selected for 3D reconstruction. These particles were reextracted at 512-pixel box size and subjected to 3D Ab-Initio reconstruction of CryoSPARC to build four initial models. A good class containing 180,924 particles were selected to execute Homogenerous Refinement. This resulted in a map at a global resolution of 3.16 Å. According to the Refinement, two flexible regions were masked for local refinement, reaching a resolution of 3.78Å and 3.24Å, respectively. By subtracting the signal of the flexible regions, a map at 3.13Å resolution were obtained through local refinement.

### Model building and refinement of the spinach PEP complex

To build an atomic model, predicted structures of subunits in *Arabidopsis thaliana* were employed from Alphafold^23,24^ and SWISS-MODEL^32^, with the atomic model of *Synechocystis* sp. PCC 6803 RNA polymerase complex (PDB: 8GZG) was fitted into density map as an initial model^27^. The initial atomic model was manually checked and mutated to fit in the map, and unidentified subunits were modeled in COOT^33^. Realspace refinement was performed in PHENIX^34^, and the COOT/PHENIX refinements were iterated until converge. Finally, all parameters were generated by PHENIX. Figures were drawn with the Pymol Molecular Graphic System (Schrödinger)^35^ and UCSF ChimeraX^36^.

## Data availability

The cryo-EM density map of the spinach PEP complex was deposited in the Electron Microscopy Data Bank (EMDB, www.ebi.ac.uk/pdbe/emdb/) under the following accession code: EMD-38799. The atomic coordinate has been deposited in the Protein Data Bank (PDB, www.rcsb.org) under the following accession code: 8XZV. All other data are available from the corresponding authors upon reasonable request.

## Acknowledgements

We are also grateful to staff of the National Facility for Protein Science (NFPS) for instrument support and technical assistance during cryo-EM data collection. This work was supported in part by the National Key R&D Program of China (No. 2020YFA0907600 and 2022YFC3401800), the National Natural Science Foundation of China (32000444 and 32000184), Shandong Provincial Natural Science Foundation (ZR2019ZD48), the Strategic Priority Research Program of CAS (XDA26050402) and the Science & Technology Specific Project in Agricultural High-tech Industrial Demonstration Area of the Yellow River Delta (2022SZX12).

## Author contributions

L.-J.Y. and C.L. conceived the project. T.W, Y.F., Y.Z., W.P., Y.Z. and A.Z. performed the sample preparation and characterization. G.-L.W. collected the cryo-EM data. G.-L.W. and L.-J.Y. processed the cryo-EM data and reconstructed the cryo-EM density map. G.-L.W. L.-J.Y. and Y.F. built the structure model and refined the structure. Y.F., G.-L.W. and Y.Z. prepared figures. G.-L.W., L.-J.Y. and Y.F. analyzed the structure. G.-L.W., Y.F., Y.Z., C.L. and L.-J.Y. jointly wrote the manuscript. All authors contributed to the discussion and comments on the results and the manuscript.

