## Extended data fig.1-9 and Extended data table 1-2 for "Architecture of the spinach plastid-encoded RNA polymerase"

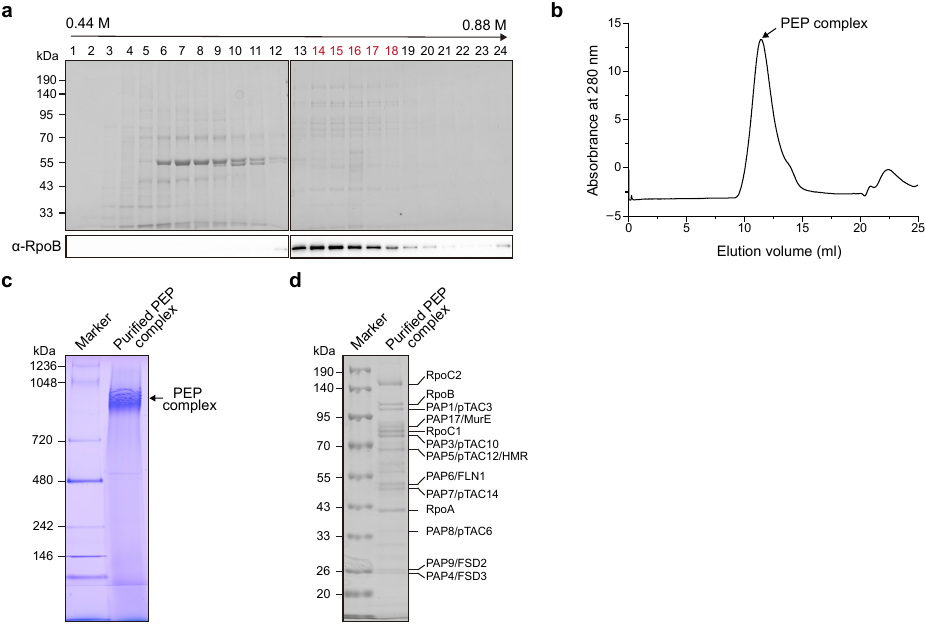

**Extended Data Fig.1 | Sample preparation and characterization of the PEP complex from spinach.** **a**, Separation of the PEP complex by sucrose density gradient (SDG) centrifugation from the preparations eluted from heparin column. After centrifugation, 24 fractions were collected from the top to the bottom of the gradient. Each fraction was resolved by electrophoresis and subjected to immunoblot analysis. The fractions (highlighted by red color) that contain PEP complex were applied for size-exclusion chromatography. **b**, Size-exclusion chromatographic elution profile of the PEP complex fractions isolated by SDG. Superose 6 Increase 10/300 GL column was used, the flow rate was set to 100 0.5 ml min^–1^, and monitored by absorption at 280 nm. **c**, The purified spinach PEP complex was separated on 4–12% blue native polyacrylamide gel electrophoresis (BN-PAGE). **d**, The purified spinach PEP complex was separated on 10% SDS-PAGE.

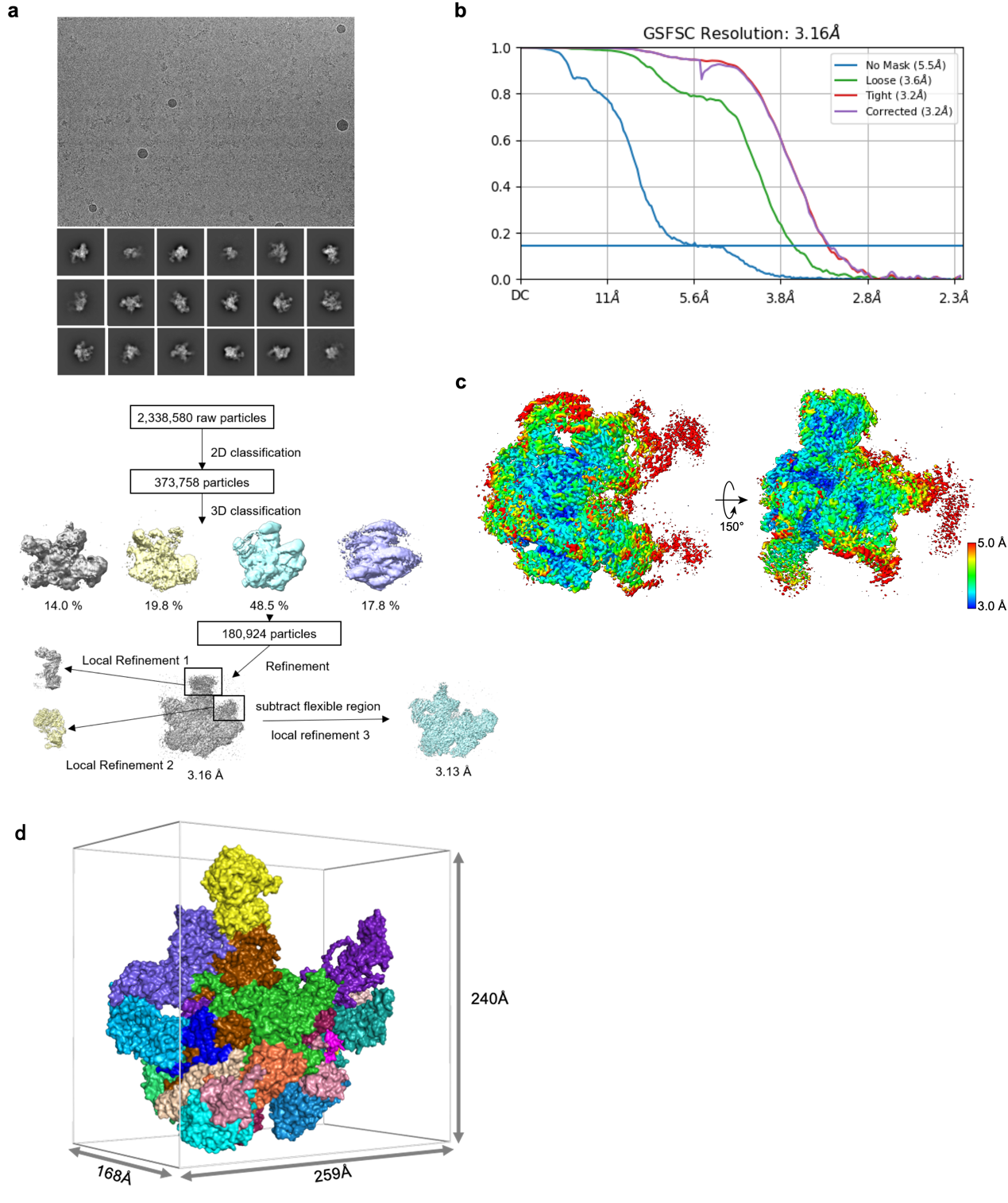

**Extended Data Fig. 2 | Data collection and image processing of PEP complex particles visualized by cryo-EM.** **a**, Flow chart of the cryo-EM data processing of the overall PEP complex and local-refinement of the PEP complex. All data were collected on a K3 direct electron detector. **b**, FSC curves of the final 3D reconstruction of the PEP complex. **c**, Local resolution distributions of the PEP complex generated with cryoSPARC. **d**, Size dimension of the spinach PEP complex.

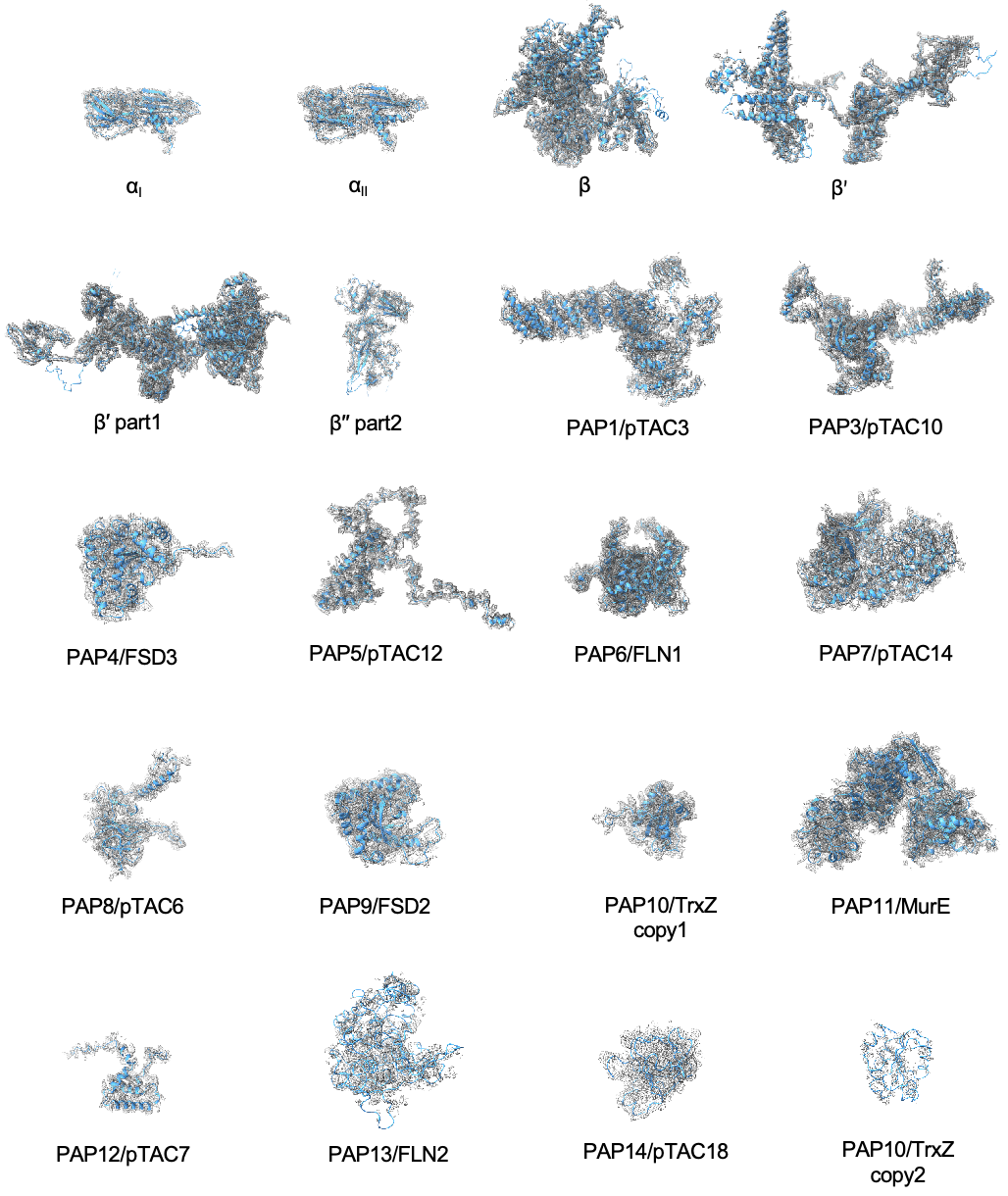

**Extended Data Fig. 3 | Cryo-EM density maps of protein subunits in the spinach PEP complex.** The cryo-EM density maps of each subunit are depicted in gray meshes. The structural models of each PEP subunit are shown as cartoon and docked into its corresponding cryo-EM densities.

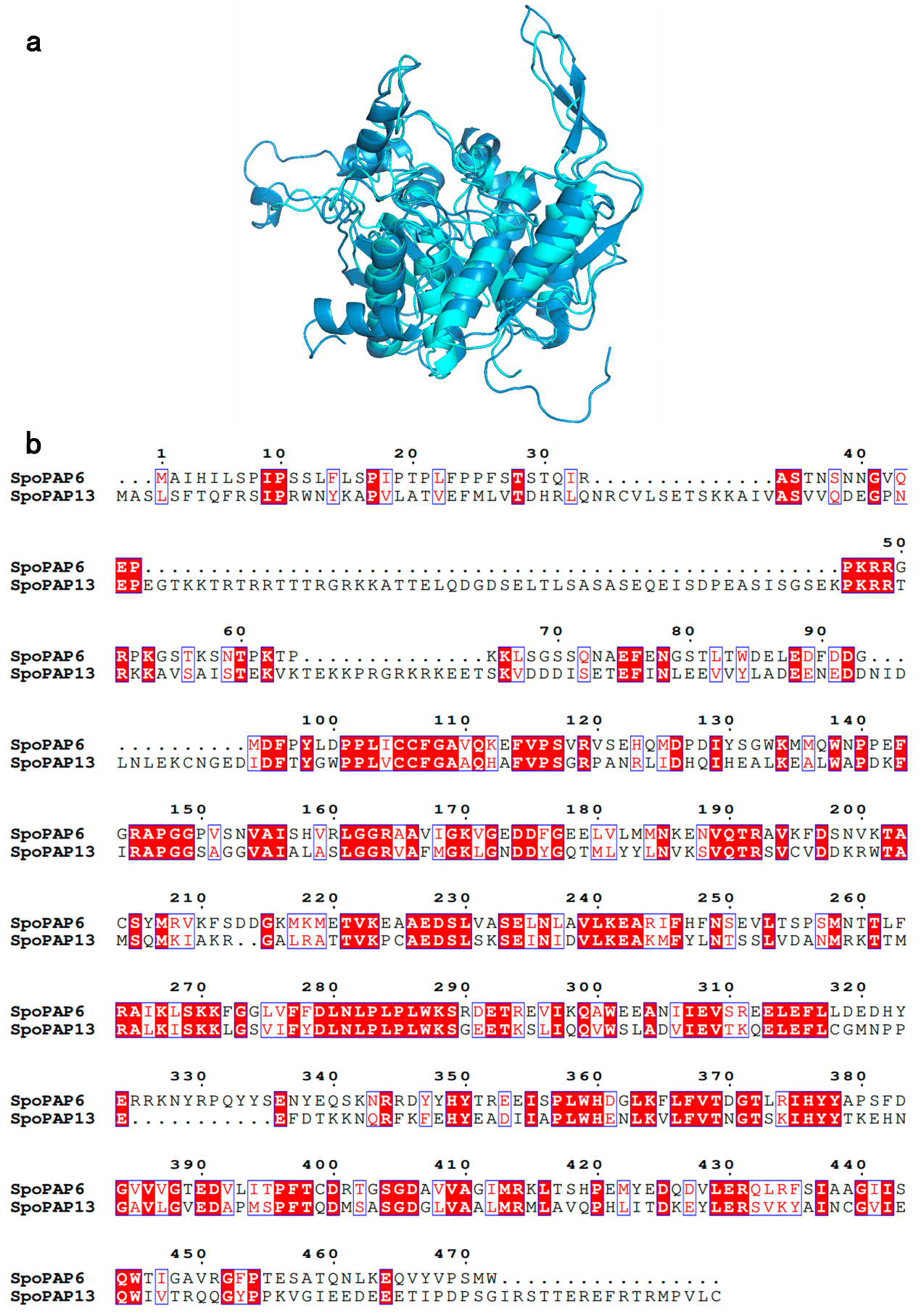

**Extended Data Fig. 4 |** The comparison of PAP6 and PAP13 from PEP complex. a, The structural comparison between PAP6 and PAP13 from PEP complex. b, The sequence alignment between PAP6 and PAP13 from PEP complex. PAP6 and PAP13 are colored individually as indicated in Figure 1.

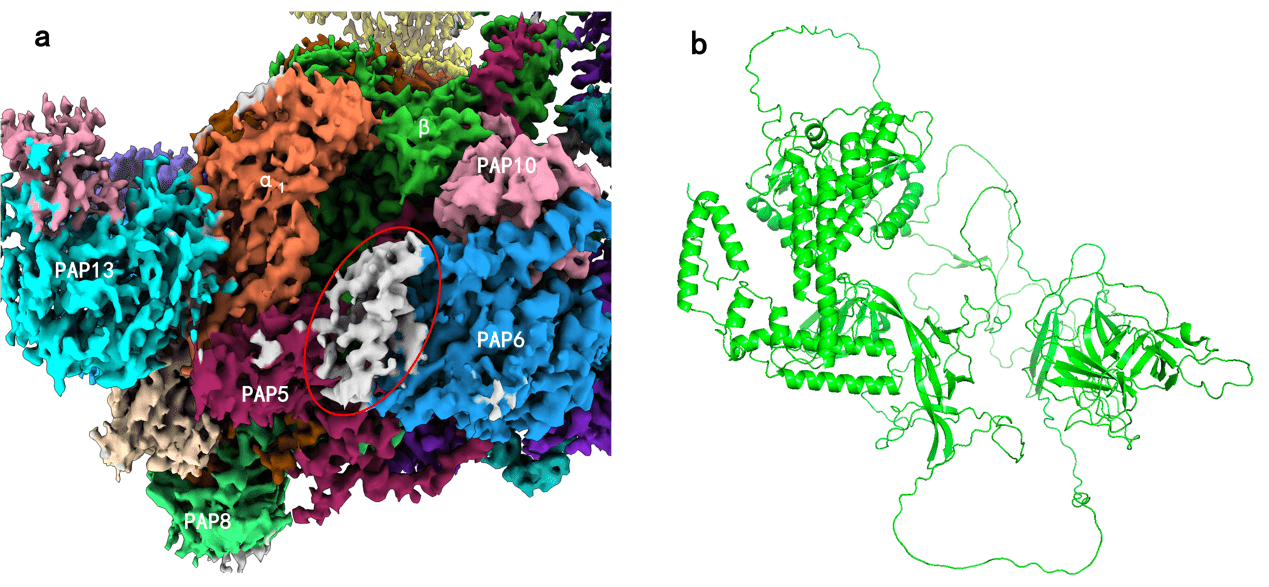

**Extended Data Fig. 5 | Uncertainties in structure determination. a,** Density map of the PEP complex. The regions involving unassigned fragmented electron densities are highlighted by a red cycle. **b**, The structure of the β'' subunit from the spinach PEP complex predicted by AlphaFold2.

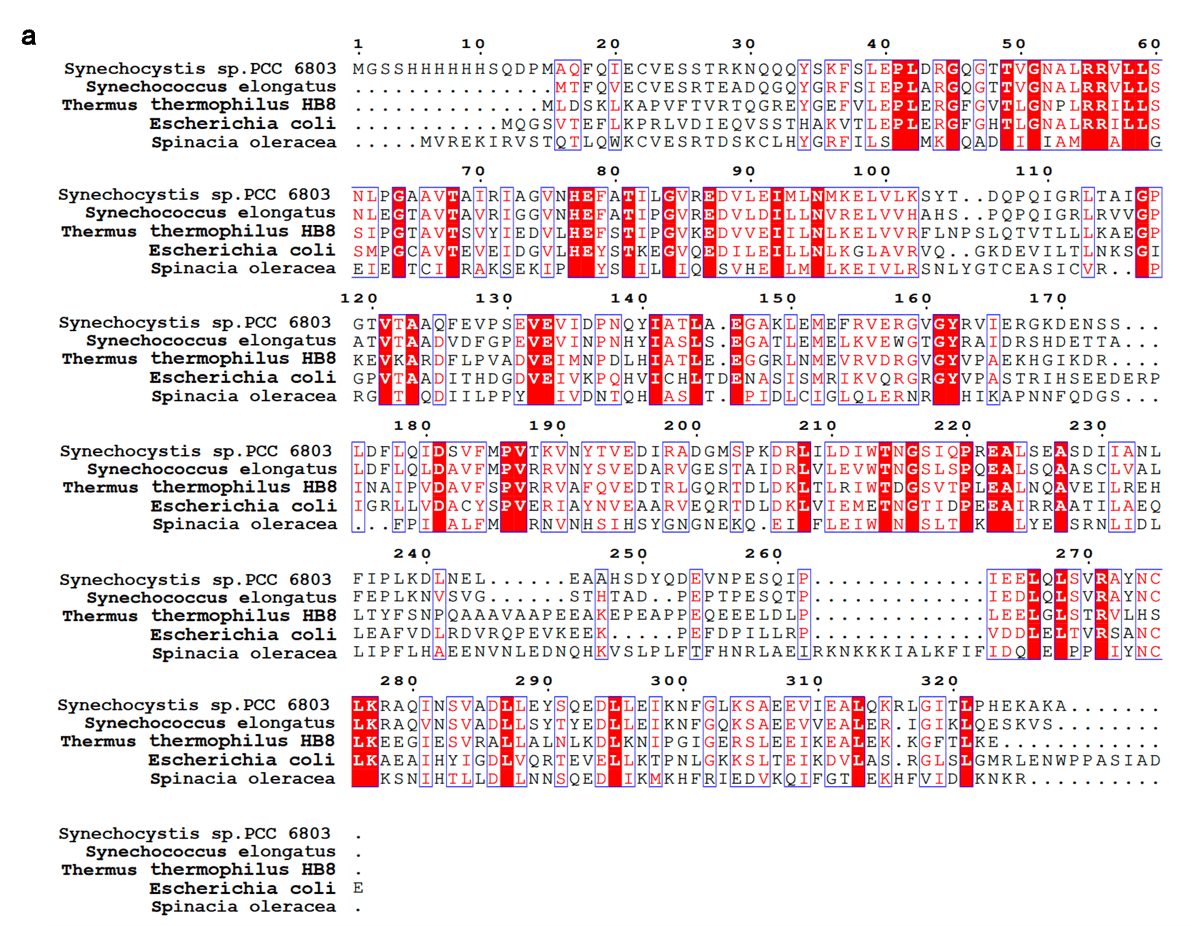

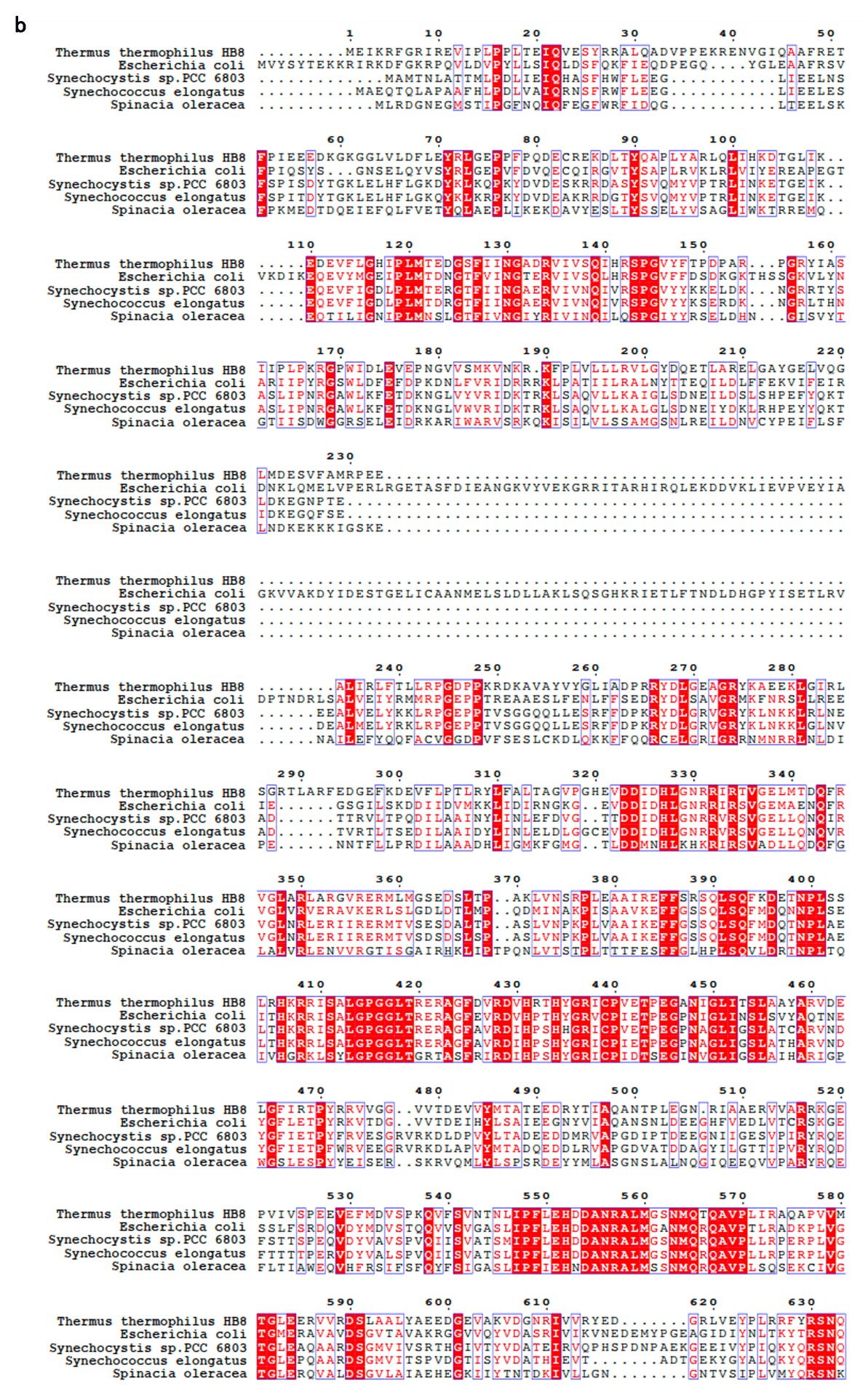

Continued on the next page

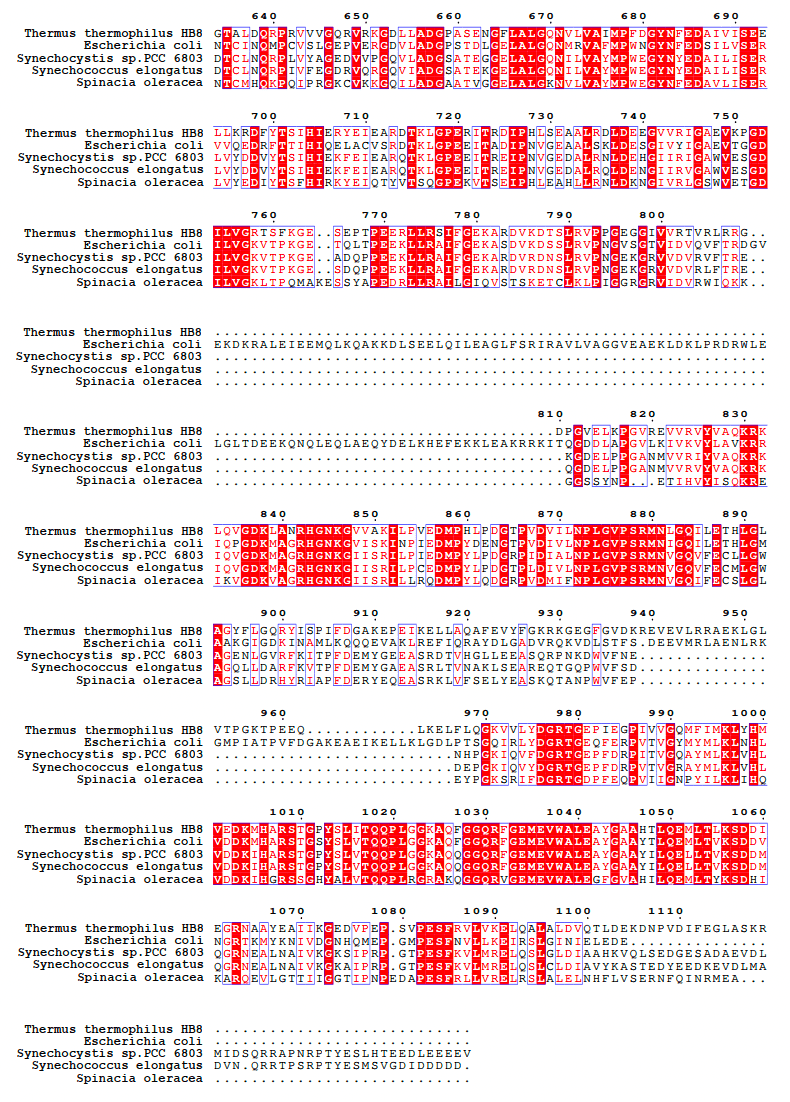

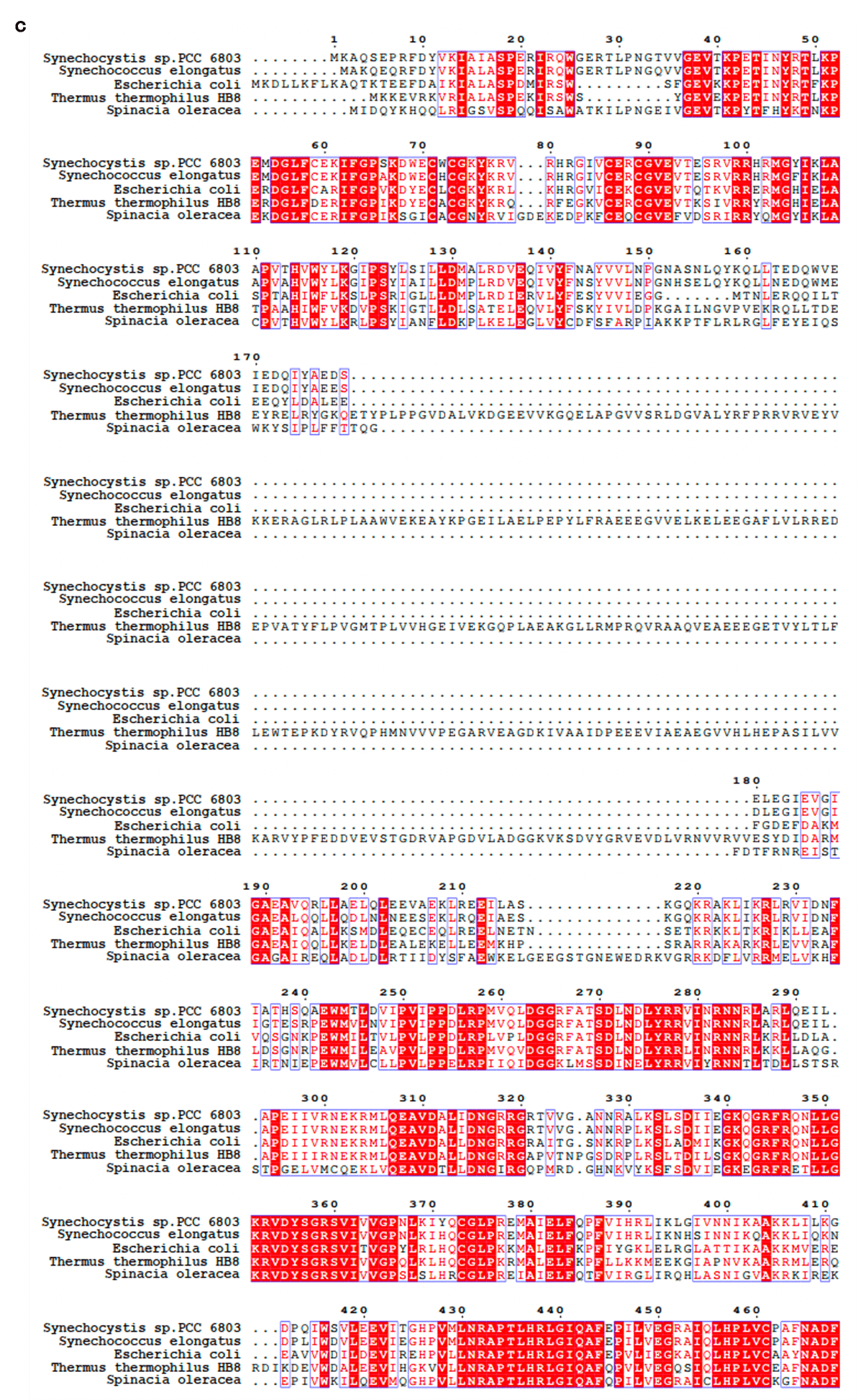

Continued on the next page

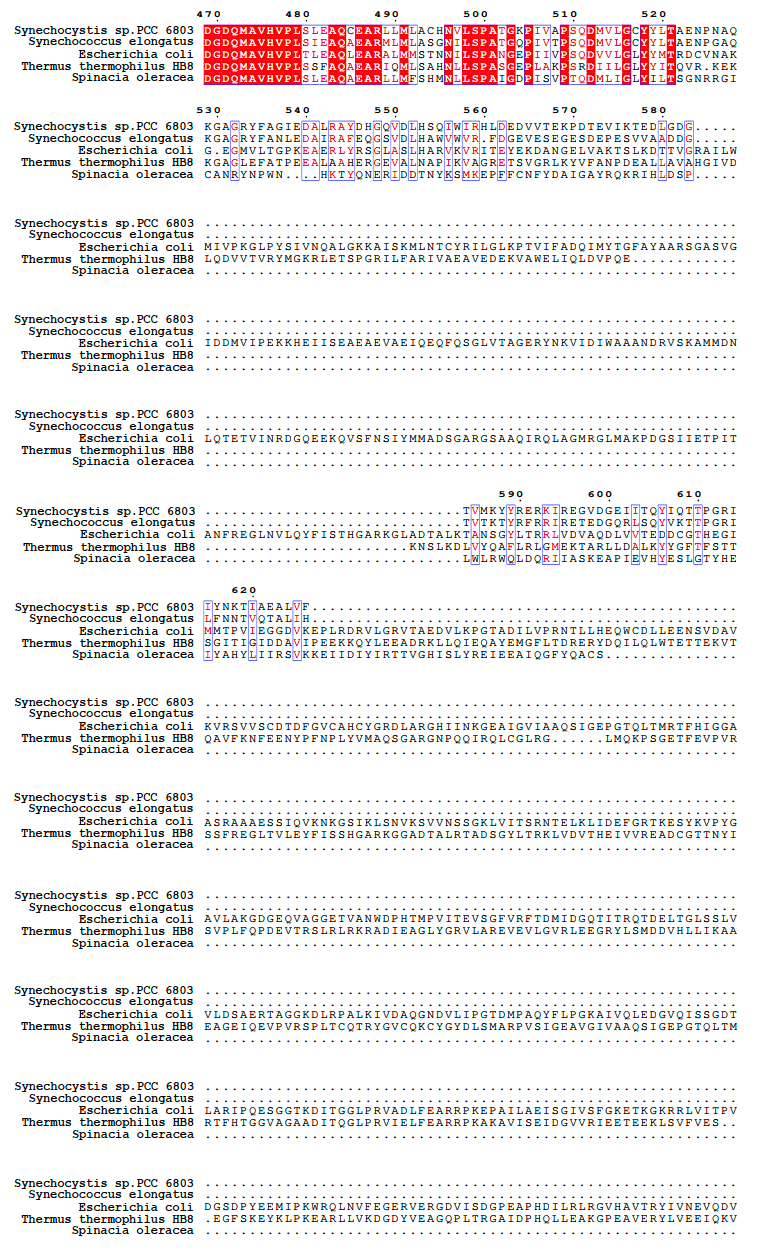

Continued on the next page

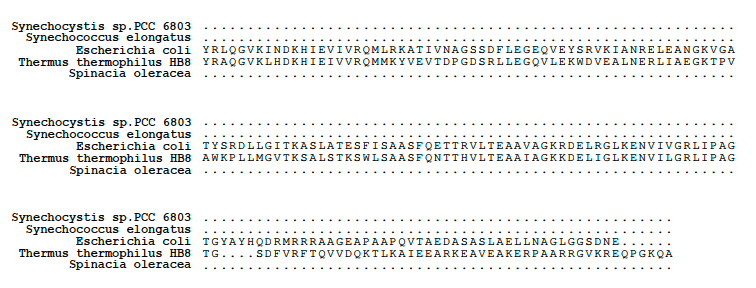

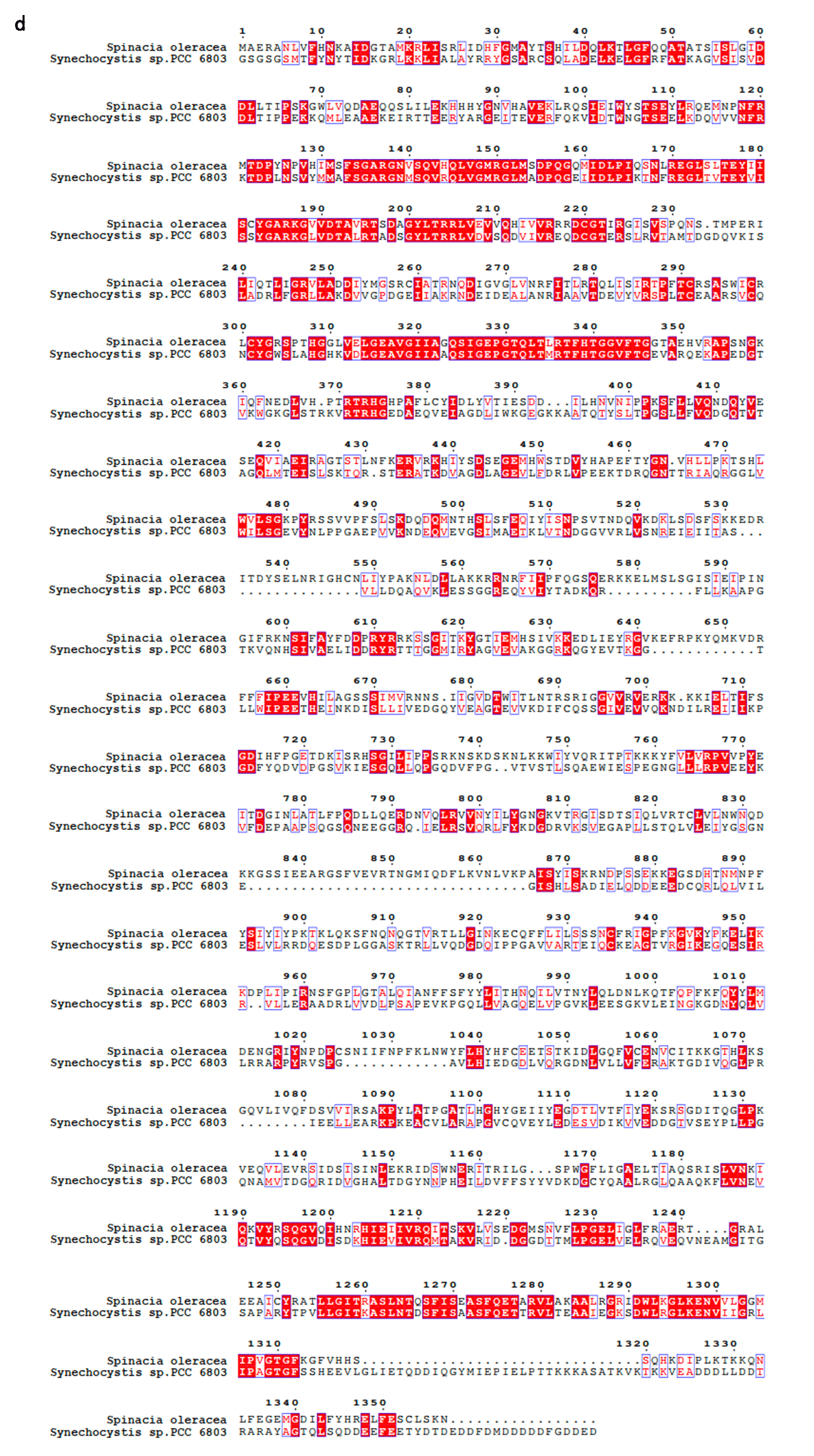

**Extended Data Fig. 6 | Multiple sequence alignments of the spinach PEP core subunits with those of from other species. a and** **b,** Sequence alignment of α, β subunit of the PEP core with α and β subunits from *Thermus thermophilus*, *E. coli*, *Synechocystis sp.* PCC 6803, *Synechococcus elongatus*. **c,** Sequence alignment of β' subunit of the PEP core with β' subunits from *Thermus thermophilus* HB8 and *E. coli* and with γ subunit from *Synechocystis sp.* PCC 6803 and *Synechococcus elongatus* (encoded by *RpoC1*). **d,** Sequence alignment of β'' subunit of the PEP core with β' subunit from *Synechocystis sp.* PCC 6803 (encoded by *RpoC2*)

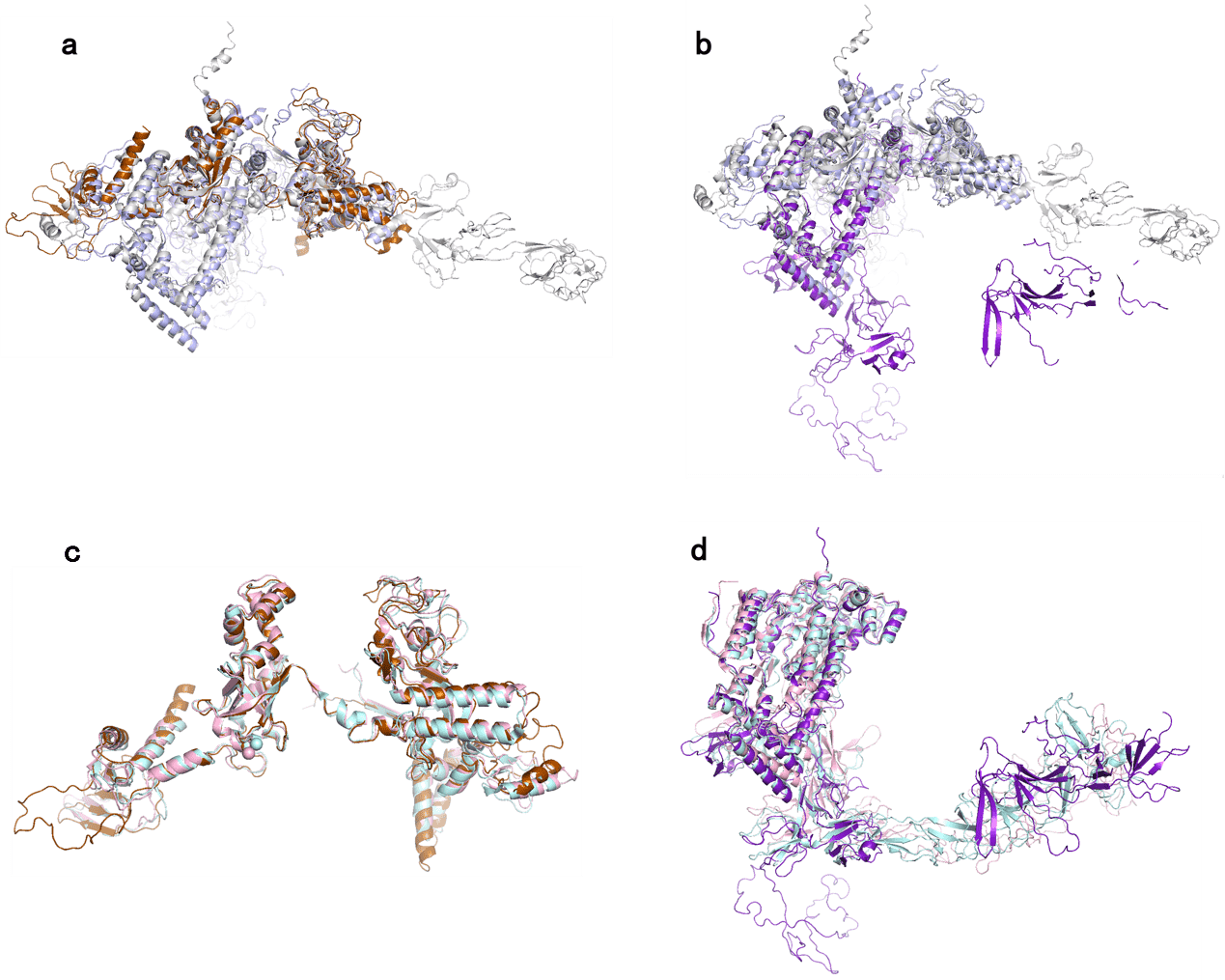

**Extended Data Fig. 7 | The structural alignments of β' and β''from the spinach PEP complex with the β' subunit of bacterial RNAP.** **a,** The superposing of the β' subunit from the spinach PEP core to the β' subunit of *E. coli* RNAP (PDB code: 6GH5; light blue) and *Thermus thermophilus RNAP* (PDB code: 3DXJ; light gray). **b**, The structural alignment of the β' subunit from the spinach PEP core to the β' subunit of *E. coli* RNAP (PDB code: 6GH5; light blue) and *Thermus thermophilus RNAP* (PDB code: 3DXJ; light gray). **c**, The structural alignment of the β' subunit from the spinach PEP core to the γ subunit from *Synechocystis sp.* PCC 6803 (PDB code: 8GZG; pale cyan) and *Synechococcus elongatus* RNAP (PDB code: 8URW; light pink). **d**, The structural alignment of the β'' subunit from the spinach PEP core to the β' subunit from *Synechocystis sp.* PCC 6803 (PDB code: 8GZG; pale cyan) and *Synechococcus elongatus* RNAP (PDB code: 8URW; light pink). The spinach PEP core subunits are colored individually as indicated in Figure 1.

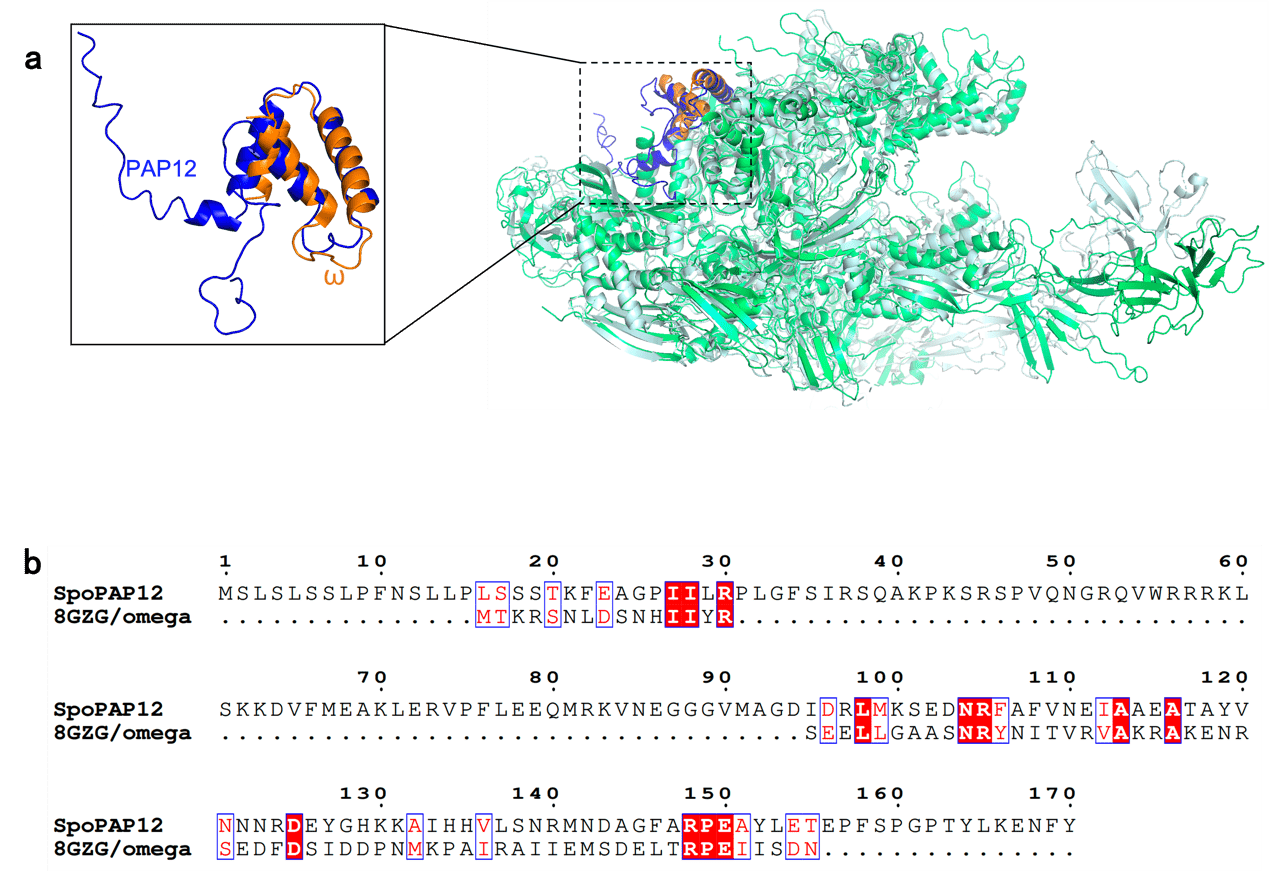

**Extended Data Fig. 8 | Comparison of PAP12 and cyanobacterial ω factor. a,** The structural alignment between the PAP12 subunit of the PEP complex and the ω subunit of *Synechocystis sp.* PCC 6803 RNAP (PDB code: 8GZG). The PEP core is colored as green. The *Synechocystis sp.* PCC 6803 RNAP is colored as light cyan. The PAP12 subunit of the spinach PEP complex is colored as blue and the ω subunit of *Synechocystis sp.* PCC 6803 RNAP is colored as orange. **b,** The sequence alignment between the PAP12 subunit of the PEP complex and the ω subunit of *Synechocystis sp.* PCC 6803 RNAP.

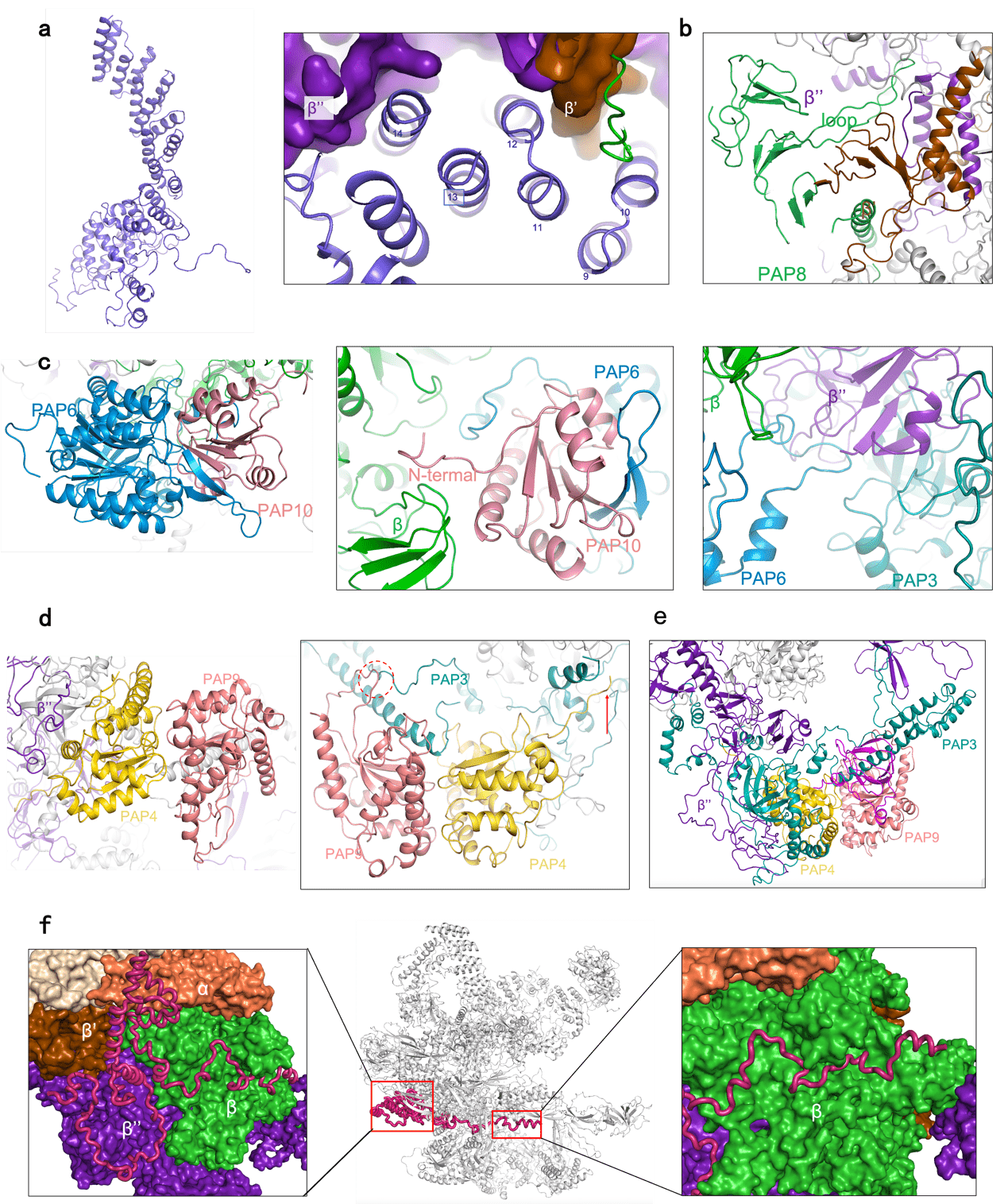

**Extended Data Fig. 9 | The interactions among different subunits in the spinach PEP complex. a**, The overall structure of the PAP1 subunit and its interaction details with the PEP core. **b**, The interaction details of the PAP8 subunit with the PEP core. **c**, The overall structure of PAP6/PAP10 heterodimer and their interaction details with the PEP core. **d**, The overall structure of PAP4/PAP9 heterodimer and their interaction details with the PAP3 subunit. **e**, The interaction details of the PAP3 subunit with the PEP core and other PAP subunits. **f**, Location and interaction details of the PAP5 subunit with the PEP core. All subunits are colored individually as indicated in Figure 1.

**Extended Data Table 1 | Cryo-EM data collection, refinement and validation statistics.**

|  | The PEP complex  (EMD-38799, PDB ID: 8XZV) |
| --- | --- |
| **Data collection and processing** |  |
| Magnification | 64000 × |
| Voltage (kV) | 300 |
| Electron exposure (e–/Å^2^) | 50 |
| Defocus range (μm) | -0.8 to -2.0 |
| Pixel size (Å) | 0.55 |
| Symmetry imposed | C1 |
| Initial particle images (no.) | 2,338,580 |
| Final particle images (no.) | 180,294 |
| Map resolution (Å) | 3.16 |
| FSC threshold | 0.143 |
| **Refinement** |  |
| Initial model used (PDB code) | 8GZG |
| Model resolution (Å) | 3.8 |
| FSC threshold | 0.5 |
| Map sharpening *B* factor (Å^2^) | -65.8 |
| Model composition |  |
| Non-hydrogen atoms | 58986 |
| Protein residues | 7499 |
| Ligands | 1 |
| *B* factors (Å^2^) |  |
| Protein | 254.84 |
| Ligand | 91.94 |
| R.m.s. deviations |  |
| Bond lengths (Å) | 0.007 |
| Bond angles (°) | 0.832 |
| Validation |  |
| MolProbity score | 2.3 |
| Clash score | 15.81 |
| Poor rotamers (%) | 0.36 |
| Ramachandran plot |  |
| Favored (%) | 88.26 |
| Allowed (%) | 11.18 |
| Disallowed (%) | 0.56 |

**Extended Data Table 2 | Subunits of the spinach PEP complex identified in the present study.**

| Protein name | Gene ID |
| --- | --- |
| PAP1/pTAC3 | 110792506 |
| PAP3/pTAC10 | 110805032 |
| PAP4/FSD3 | 110793156 |
| PAP5/pTAC12 | 110799664 |
| PAP6/FLN1 | 110803951 |
| PAP7/pTAC14 | 110796283 |
| PAP8/pTAC6 | 110782000 |
| PAP9/FSD2 | 110788339 |
| PAP10/TrxZ | 110801385 |
| PAP11/MurE | 110791626 |
| PAP12/pTAC7 | 110783140 |
| PAP13/FLN2 | 110797280 |
| PAP14/pTAC18 | 110798943 |
| RpoA/α | 2715631 |
| RpoB/β | 2715632 |
| RpoC1/β' | 2715633 |
| RpoC2/β'' | 2715634 |
